# Transcriptional profiling defines unique subtypes of transit amplifying neural progenitors within the neonatal mouse subventricular zone

**DOI:** 10.1101/2025.08.20.669012

**Authors:** Rebecca Zaritsky, Ekta Kumari, Fernando Janczur Velloso, Alexander Lemenze, Seema Hussain, Steven W. Levison

## Abstract

While significant progress has been made in understanding the heterogeneity of Neural Stem Cells (NSCs), our understanding of similar heterogeneity among the more abundant transit amplifying progenitors is lagging. Our work on the neural progenitors (NPs) of the neonatal subventricular zone (SVZ) began over a decade ago, when we used antibodies to the 4 antigens, CD133, LeX, CD140a, and NG2 to perform FACS to classify subsets of the neonatal SVZ as either multi-potential (MP1, MP2, MP3, MP4 and PFMPs), glial-restricted (GRP1, GRP2, and GRP3) or neuron-astrocyte restricted (BNAP). Using RNASeq, we have characterized the distinctive molecular fingerprints of 4 SVZ neural progenitors and compared their gene expression profiles to those of the NSCs. We performed bioinformatic analyses to provide insights into each NP type’s unique interactome and the transcription factors regulating their development. Overall, we identified 1581 genes upregulated in at least one NP compared to the NSCs. Of these genes, 796 genes were upregulated in BNAP/GRP1 compared to NSCs; 653 in GRP2/MP3; 440 in GRP3; and 527 in PFMPs. One gene that emerged from our analysis that can be used to distinguish the NPs from the NSCs is Etv1, also known as Er81. Also notable is that the neural stem cells downregulated cilia formation genes as they differentiated to become multipotential progenitors. Among the NPs, both PFMP and GRP3 subtypes differentially expressed genes related to neuron and oligodendrocyte development, including Matn4, Lhfpl3 and Olig2. GRP3s uniquely expressed Etv5, a transcription factor known to promote glial cell fate specification, while PFMPs uniquely expressed Lhx6, a transcription factor that regulates interneuron specification. PFMPs also expressed transcripts for olfactory receptors. Unlike the other NPs, the GRP1 and GRP2 NPs upregulated expression of genes for proteins involved in immune function. The present work will serve as an important resource for investigators interested in further defining the transit amplifying progenitors of the mammalian SVZ.

## 1. Introduction

Mammalian neural development is a dynamic process that begins two weeks into pregnancy and progresses into adolescence. In early pregnancy, pyramidal neurons and astrocytes are generated from the ventricular zone (VZ). As neural development progresses, the VZ regresses and the subventricular zone (SVZ) expands to generate the bulk of the brain’s interneurons and glia. There is increased interest in studying the SVZ as it remains active throughout the lifespan of rodents and through the first years of life in humans. It can potentially be stimulated or reprogrammed neurogenesis to generate new neurons or glia to treat brain injury.

The subventricular zone of the adult has been extensively characterized. Neural stem cells (NSCs), often referred to as “Type B cells” [1,2], have ciliated processes that contact the cerebrospinal fluid, enabling them to respond to growth factors and cytokines that maintain their primitive state [3]. NSCs arise in the SVZ during mid-embryonic development [4] and divide symmetrically [5] to produce intermediate progenitors (Transit Amplifying Progenitors; TAPs) [6], originally called “Type C cells”, which then become lineage-restricted to form terminally differentiated cells [6]. SVZ-born TAPs predominantly give rise to interneurons in the olfactory bulb and striatum, as well as to astrocytes and oligodendrocytes in the cortex, striatum, and white matter [7-9].

Increasingly, there is interest in characterizing the NSCs and TAPS of the fetal and neonatal brain, as SVZ-born progenitors are critical for proper neural development. Notably, the net number of neurons in the cerebral cortex doubles during the first two weeks of rodent postnatal life and the majority of glial cells are born postnatally [10,11]. Subsequently, current evidence indicates that deficits in NSC proliferation and differentiation represent the major convergence point for neurodevelopmental disorders (NDDs), and mutations in genes that play a role in this process are implicated in the majority of NDDs [12,13]. Correspondingly, postnatal neurogenesis is profoundly impaired in NDDs, and manipulating neurogenesis can ameliorate phenotypes [14].

There has been much debate in the neuroscience community about the definition of a “stem cell” vs an “intermediate progenitor” [15]. Many studies have characterized cells as NSCs based on their ability to form neurospheres or because they are tripotential, which is inadequate given that many intermediate progenitors form neurospheres and are tripotential. However, unlike NSCs, intermediate progenitor cells do not exhibit self-renewal after several generations [16]. Therefore, understanding the differences between NSCs and the variety of neural progenitor subtypes is crucial to understand the transitions that occur during neurodevelopment.

Previous studies that aimed to investigate the diversity of cells generated postnatally in the SVZ have used scRNAseq on unsorted [17-19] and FACS-sorted cells [20,21], and used bioinformatics to classify the cells as NSCs, epithelial, neuronal, glial and immune lineages. Although these seminal studies describe some subtypes of TAPs, the emphasis of their analyses was to characterize the NSC subpopulations and terminally differentiated cells. Moreover, these studies classified adult SVZ cells where the variety of cell types being produced is restricted. Thus, the diversity and function of TAPs during early postnatal neurodevelopment in mice remain largely uncharacterized.

In 2007, David Panchision and co-workers combined antibodies against CD133, CD15, CD24, A2B5 and PSA-NCAM, to identify and enrich for 4 sets of neural precursors from the E13.5 and P2 mouse VZ/SVZ. They established that there were multipotent progenitors that could produce neurons, astrocytes and oligodendrocytes, that there were progenitors that were bipotential and produced either neurons and oligodendrocytes, or astrocytes and oligodendrocytes, and that there were progenitors that only produced neurons [22]. Extending Panchision’s studies, we combined CD133 and LeX and then added two intermediate progenitor antigens, CD140a and NG2 [23,24]. With this strategy, 8 phenotypically defined subsets of neural progenitors could be identified within the periventricular region [25]. To determine the developmental potentials of these 8 subpopulations, SVZ neurospheres were generated, dissociated into a single cell suspension, separated by FACS, plated onto laminin-coated chamber slides at low density, and expanded with growth factors. The multipotential progenitors (MPs), included the NSCs, MP1, MP2, MP3, MP4 and the platelet-derived growth factor-fibroblast growth factor responsive MP cell (PFMP). There were 4 types of bipotential progenitors identified that included the bipotential neuronal-astrocytic progenitor (BNAP) and 3 glial-restricted progenitors (GRP1, GRP2 and GRP3). Building upon that work, here we have isolated the four largest populations of progenitors (BNAP/GRP1, GRP2/MP3, GRP3, and PFMP) using FACS and then subjected those cells to RNA sequencing. This approach allows us to analyze the transcriptome of early postnatal SVZ progenitors characterized by established antigenic profiles. In doing this, we can illuminate the stages of neurodevelopment from NSCs to mature neurons and macroglia.

## 2. Materials and Methods

### 2.1. Mice

Swiss Webster Mice from timed-pregnant females purchased from Charles River laboratories or generated in house were used for all studies. Each litter was analyzed collectively with both males and females. All experiments were performed in accordance with Protocol #999901108 approved by the IACUC committee of Rutgers University Biomedical Health Sciences.

### 2.2. SVZ Cell Dissociation

Mice were terminated via decapitation at postnatal day 5. The periventricular zone was micro-dissected from coronal sections on ice and into Flow Buffer (DMEM/F12 Media with 20mM HEPES, 1% BSA, and 1mM EDTA, pH 7.4). All pups from each litter were pooled for further experiments. Cells were minced then incubated in an enzyme solution (DMEM/F12 with 0.1Wu Liberase-DH, 10ug/mL DNase I, and 20mM HEPES) for 15 minutes with rocking at 37°C. 1mL FBS and 4mL of Flow Buffer was added and cells were centrifuged for 5 minutes at 300g. Cells were manually triturated using a P1000 tip until no visible debris remained, triturated using a 200uL pipette tip, and filtered through a 30 micron cell filter. Cells were then washed in a polystyrene tube to remove dead cells and debris.

### 2.3. Flow Cytometry and Cell Sorting

Flow cytometry was performed as described previously by our group [25-27] using the following antibodies: Lewis-X (1/20; BD Biosciences), CD133-APC (1/50; eBioscience), CD140a (1/400; BioLegend), and NG2 chondroitin sulfate proteoglycan (1/50; Millipore). Goat anti-rabbit IgG Alexa Fluor 700 (1/100; Invitrogen) was used for NG2 and DAPI (1/50,000) was used for dead cell exclusion. Cells were collected into polypropylene tubes of lysis buffer using the FACS BD Aria II.

### 2.4. RNA Sequencing

RNA was isolated from SVZ neural progenitors (NP) using NucleoSpin® RNA XS from Macherey Nagel (Germany) according to the manufacturer’s instructions. The integrity of the RNA was analyzed on Bioanalyzer 2100 using an RNA Pico6000 Kit (Agilent Technologies). All RNA samples used in this study had an RNA integrity number (RIN) >7.0. PolyA plus RNAs were amplified and enriched using Ambion’s messageAmp II aRNA amplification kit. The quality of amplified aRNA was verified using the Bioanalyzer. A cDNA library was constructed using the Illumina TrueSeq protocol.

The barcoded cDNA library was purified with AmpureXP beads and quantified using Qubit High sensitivity DNA kit and the quality of the library was determined by analyzing on Agilent bioanalyzer. Libraries mixed at equimolar ratios were sequenced on a NextSeq500 (Illumina, San Diego, CA) for 75 cycles.

### 2.5. Bioinformatics Analysis

Data were processed using nf-core/rnaseq v3.10 (doi: https://doi.org/10.5281/zenodo.1400710) of the nfcore collection of workflows [28] and executed with nextflow v22.10.11. Briefly, raw transcriptome reads were assessed for quality control (FASTQC v0.11.9) and trimmed for quality and adapter contaminants (cutadapt v3.4). Trimmed reads were aligned to the Mus musculus genome GRCm38 using STAR (v2.6.1), followed by transcript abundance calculation via Salmon (v1.9.0). 2 experiments were excluded because the samples had low quality, and 4 experiments were analyzed further. Hit count normalization and differential gene expression group cross-comparisons were performed using DESeq2 (v1.26.0). Significantly differentially expressed gene thresholds were set at FDR-adjusted p <0.05. Up- and down-regulated gene lists were generated for each NP subtype and composed of all genes with differential expression compared to NSCs (p < 0.05 adjusted). Each gene list was analyzed using Network Analyst to identify protein-protein interactions and transcription factor networks. Overrepresentation analysis was performed using the clusterProfiler package in R (version 4.16.0) to identify up- and down-regulated gene ontology terms in each population compared to NSCs.

### 2.6. NSC Upset Plot Generation

An Upset plot comparing NSC-enriched genes was generated using UpsetR v1.4.0 in R. To identify genes upregulated in NSCs in our dataset, we used DESeq2 to compare NSC transcripts to the transcripts expressed collectively in all of the NPs. Transcripts with higher expression in NSCs than NPs (log2 Fold change > 0.25 and p < 0.05) were classified as NSC-enriched. We compared our dataset with the publicly available data from the articles discussed in the follow paragraph. The code used to analyze data and generate plots is available on our Github Repository.

Cebrian-Silla et al., (2021) [29], Zywitza et al., (2018) [17], Dulken et al., (2017) [20] and Beckervordersandforth et al., (2010) [21] analyzed the subventricular zone of adult mice. Marcy et al., (2023) [30] used P12 mice and Azim et al., (2015) [31] examined mice at ages P4, P8, and P11. Cebrian-Silla et al., Zywitza et al., Dulken et al., and Marcy et al. used single cell sequencing while Beckervordersandforth et al. and Azim et al. used microarrays. The datasets provided used different criteria to define a “NSC.” 3 papers used transgenic animals expressing GFP either coupled to the human GFAP (hGFAP) promoter [21,29] Hes5 [31] and all used other markers in their gene expression datasets including Prom1, Glast and Tbhs4.

Marcy et al., (2023) [30] defined NSCs based on their expression of Glast and Prom1 and non-expression of Egfr and Foxj1 (markers of activation and ependymal cells, respectively). The Seurat object was accessed using GEO Accession Number GSE186003 and analyzed using FindMarkers in R for genes enriched in qNSCs (with a threshold of Log2FoldChange > 0.25 that were expressed in >25% of NSCs with an enrichment p < 0.05 adjusted).

Cebrian-Silla et al., (2021) [29] used a hGFAP:GFP mouse and differentiated between “B cells”, parenchymal astrocytes and ependymal cells using other marker genes. The Seurat object was accessed using GEO Accession Number GSE165554 and analyzed using FindMarkers in R for genes enriched in “B cells” with a threshold of Log2FoldChange > 0.25 that were expressed in >25% of NSCs with an enrichment p < 0.05 adjusted.

Dulken et al., (2017) [20] used a hGFAP-GFP mouse and stained cells for expression of Prom1 and EGFR. Quiescent NSCs were defined as hGFAP-GFP+, Prom+ and EGFR-. The count matrix for the single cell sequencing was available in Supplementary Table S2 and was loaded into R and used to generate a Seurat object. The Seurat object was scaled and normalized and neurosphere-derived cells were excluded. The Seurat object included cells defined as qNSC, aNSC, neural progenitors and astrocytes. Genes enriched in qNSCs were calculated using FindMarkers with a threshold of Log2FoldChange > 0.25 and expressed in >25% of qNSCs with an enrichment p < 0.05 adjusted.

Beckervordsanforth et al., (2010) [21] also used a hGFAP-GFP mouse. Quiescent NSCs were defined as hGFAP-GFP+ and Prom+. Azim et al., (2015) [31] used a Hes5-GFP-expressing mouse and identified NSCs as Hes5+ and Prom1+. Zywitza et al., (2018) [17] did not use specific markers and clustered the cells before examining expression of marker genes (Thbs4, Slc1a3) to identify a NSC-cluster. Data from the Beckervordsanforth et al. Supplementary Table S6; Zywitza et al., Supplementary Table S3; and Azim et al., Supplementary Table S2 were accessed and analyzed without further modification.

## 3. Results

### 3.1. The cells of the subventricular zone differentially express surface antigens that identify unique subpopulations of neural progenitors

Cells from the mouse P5 periventricular zone were dissociated using Liberase and manual trituration. This approach preserves surface antigens that are removed using other enzymes [32,33]. The cells were stained for CD133/Prominin1, CD15/Lex/heat-stable antigen, CD140a/PDGFRα, and NG2/Chondroitin Sulfate and FACS-sorted according to antigenic profiles into five unique populations as described below. Single cells were identified using forward vs side-scatter to exclude debris and doublets, as shown in Figures 1A-1C. Dead cells were excluded using DAPI staining (Figure 1D). Live cells were then gated first according to CD140a and NG2 expression (Figure 1E) followed by a secondary gate of CD133 and CD15 (Figure 1F-1H). Neural Stem cells (NSCs), BNAP/GRP1s, GRP2/MP3s, GRP3s, and PFMPs were sorted according to antigen expression as described previously [25,27] and collected in lysis buffer for RNA extraction and further analysis.

**Figure 1:**
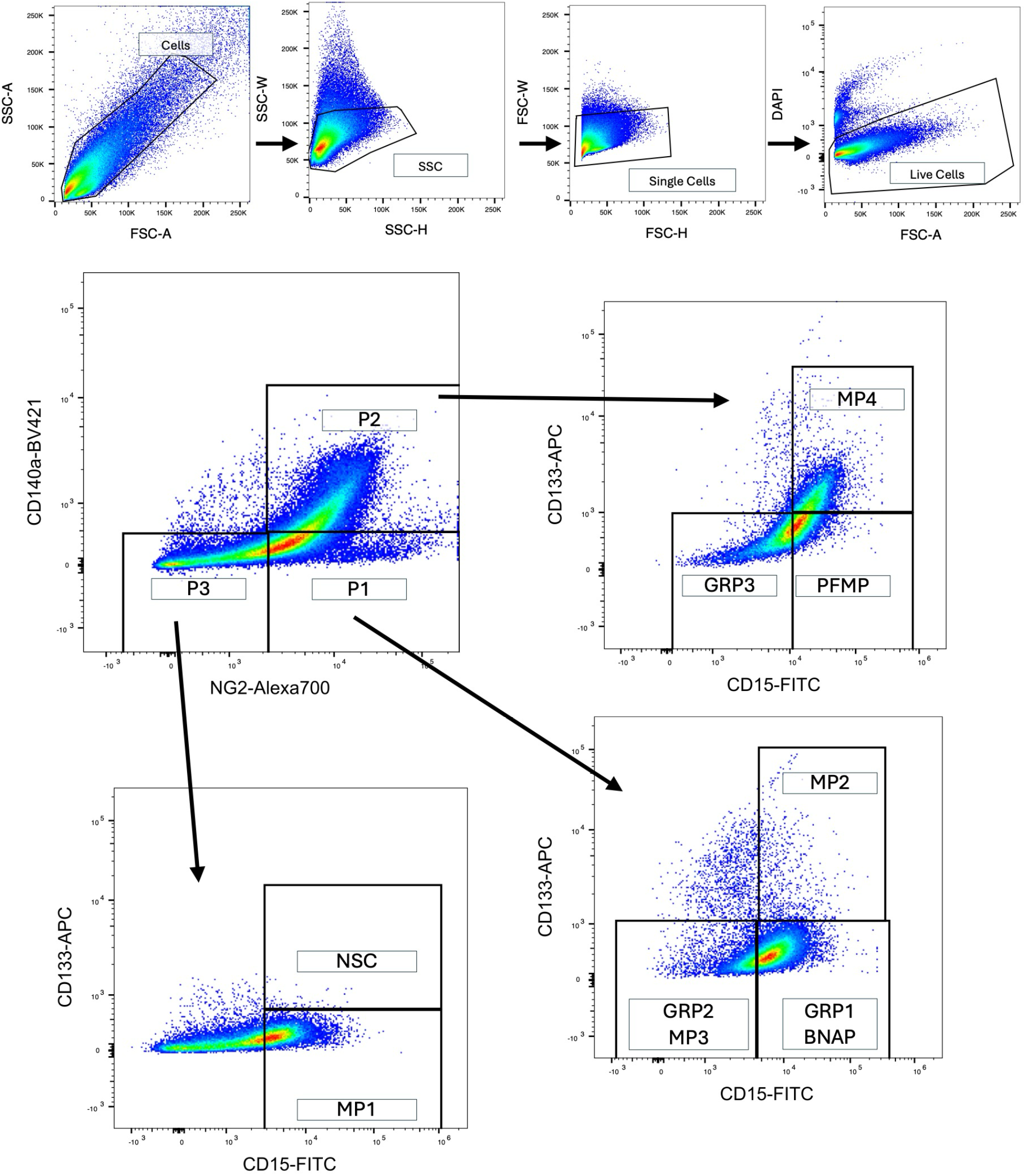
Flow cytometric analysis of the subventricular zone using a 4-color antigen panel identifies 8 unique populations of neural progenitors. The periventricular region of P5 mouse pups was dissected and cells were dissociated into a single-cell suspension using Liberase and Accutase followed by trituration. **A-C)** Debris and doublets were excluded using forward and side scatter gating and **D)** dead cells were excluded using DAPI staining. **E)** Cells were first gated on CD140a and NG2 expression followed by a secondary gate of CD133 and CD15 (**F-H)**. Neural Stem cells, BNAP/GRP1s, GRP2s, GRP3s, and PFMPs as identified in panels **F-H** were sorted and collected in lysis buffer for RNA extraction and further analysis.

### 3.2. Neural Progenitor populations have unique gene expression

RNA sequencing was performed on each of the four cell populations and on the NSCs. As each of the intermediate progenitor cells is a descendant of the NSCs, we sought to understand the genetic differences between each progenitor subtype and NSCs to understand the steps that occur in development. Therefore, subsequent analysis compared gene expression changes between NCSs and each progenitor cell type.

Overall, we identified 4647 genes with differential expression between at least one neural progenitor (NP) subtype and NSCs, of which 1581 are upregulated in at least one NP (Figure 2A), 3079 are downregulated (Figure 2B), and 13 have variable directionality between populations. 796 genes were upregulated in BNAP/GRP1 compared to NSCs; 653 in GRP2/MP3; 440 in GRP3; and 527 in PFMPs (Figure 2A). 1390 genes were downregulated in BNAP/GRP1 compared to NSCs; 1759 in GRP2/MP3; 1588 in GRP3; and 2149 in PFMPs (Figure 2B).

**Figure 2:**
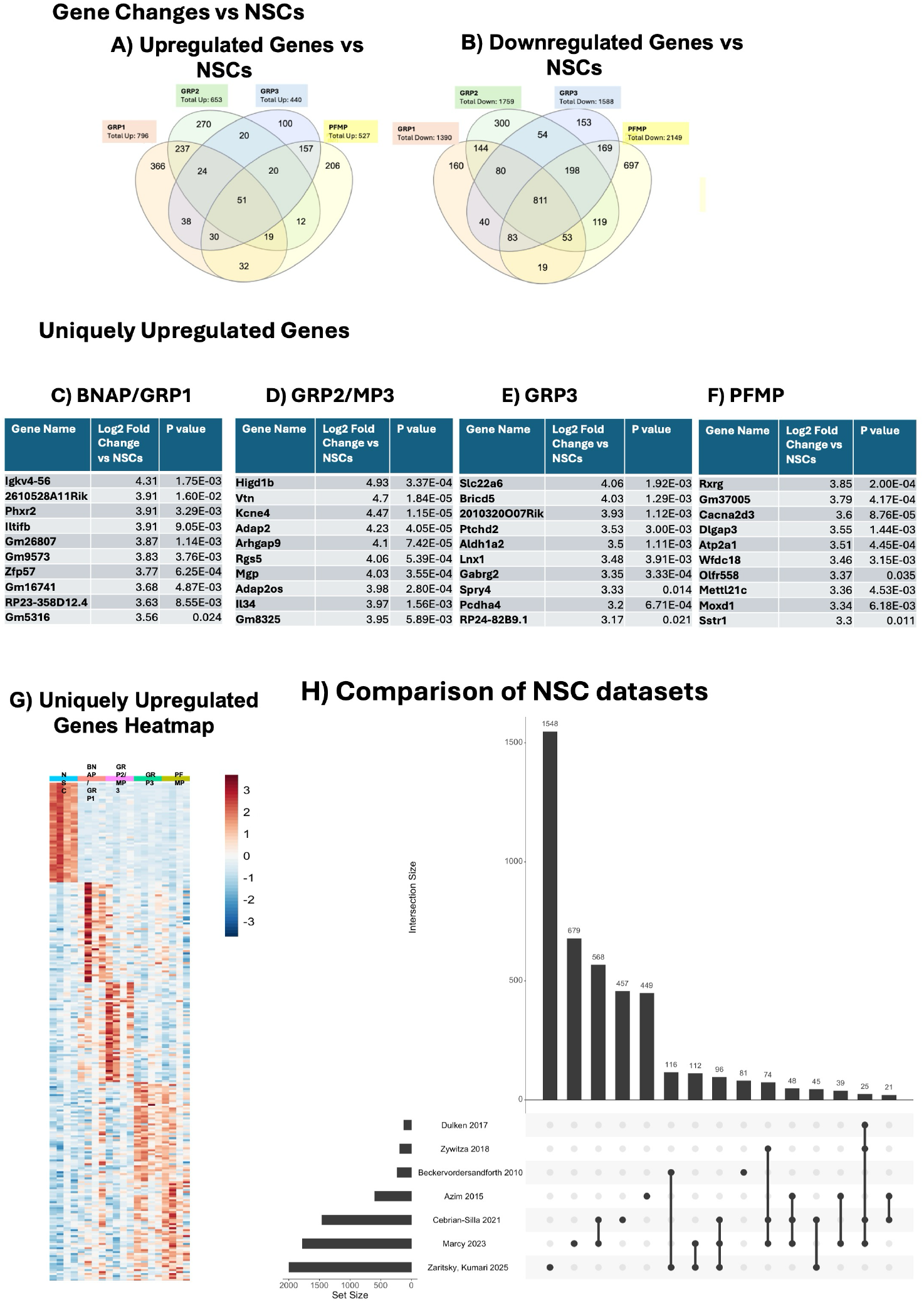
Neural progenitor subtypes are genetically distinct from neural stem cells and from each other. Neural progenitors were isolated using FACS. RNA was isolated, amplified, and analyzed using NextSeq500. Each neural progenitor population (BNAP/GRP1, GRP2/MP3, GRP3, PFMP) was compared to neural stem cells (n = 4 independent experiments). **A)** Number of genes upregulated and **B)** Number of genes downregulated in each progenitor compared to NSCs (padj < 0.05). **C-F)** Top 10 uniquely upregulated genes in each population (padj < 0.05). **C:** BNAP/GRP1; **D:** GRP2, **E:** GRP3, **F:** PFMP compared to neural stem cells **G)** Heatmap clustering the top 50 unique genes upregulated in each population. Each column represents an independent experimental replicate. **F)** Upset plot showing comparison of NSC-enriched genes identified in this paper to NSC-enriched genes from previous studies on adult and adolescent NSCs.

### 3.3. Surface markers that define NP populations correlate with gene expression

As expected by surface antigen expression, expression of the NSC marker Prom1 was down-regulated in all progenitor cell populations (p < 0.01 for all) compared to NSCs. Similarly, Pdgfra was upregulated in GRP3 and PFMP (p = 6.5E-9 and p=6.5E-8, respectively) compared to NSCs, but Pdgfra was not increased in BNAP/GRP1 and GRP2/MP3 (p = 0.27 and p =1) compared to NSCs. Interestingly, Cspg4 expression was upregulated in GRP3 and PFMP compared to NSCs (GRP3 Fold Change: 9.64, p = 4.2E-3; PFMP Fold Change: 8.37, p = 6.4E-3) but not in BNAP/GRP1 and GRP2/MP3 (BNAP/GRP1 Fold Change: 0.90, p = 1; GRP2/MP3 Fold Change: 2.23, p = 0.78).

### 3.4. Each neural progenitor population upregulates unique and shared genes compared to neural stem cells

Each NP population expressed, on average, 235 (100-366) uniquely upregulated genes compared to NSCs (Fig. 2A). The top 3 upregulated genes in each population were Igkv4-56 (fold change vs NSCs of 19.8), Cryba4 (fold change vs NSCs 19.2), and Gpr34 (fold change vs NSCs 18.7) for BNAP/GRP1; Gpr34 (fold change vs NSCs of 35.8), Gm29291 (fold change vs NSCs 32.2), and Higd1b (fold change vs NSCs 30.5) for GRP2/MP3; Lhfpl3 (fold change vs NSCs of 72.3), Dmrtb1 (fold change vs NSCs 45.9), and Matn4 (fold change vs NSCs 42.0) for GRP3; and Matn4 (fold change vs NSCs of 49.8), Tmem255b (fold change vs NSCs 49.4), and Brinp3 (fold change vs NSCs 46.7) for PFMPs (Fig. 2C-F).

### 3.5. BNAP/GRP1 progenitors share more gene expression similarities with GRP2/MP3, while GRP3s are more similar to PFMPs

There was significant overlap between upregulated genes vs NCSs in the progenitor populations, particularly between BNAP/GRP1 and GRP2/MP3 and between GRP3 and PFMP (Fig.2A). 16 of the top 50 upregulated genes were shared between BNAP/GRP1 and GRP2/MP3. For GRP3 and PFMP, 29 of the top 50 genes were shared. Overall, 237 upregulated genes were shared between BNAP/GRP1 and GRP2 and 157 between GRP3 and PFMP, while all other combinations of two or three NPs share ≤38 commonly upregulated genes. Fifty-one genes were commonly upregulated in all NPs vs NSCs, of which the highest magnitude were Igf2bp3 (8.3× in BNAP/GRP1, 11.4× in GRP2/MP3, 8.7× in GRP3, and 3.9× in PFMP); Gpr155 (6.7× in BNAP/GRP1, 5.4× in GRP2/MP3, 4.5x in GRP3, and 4.5× in PFMP); and Gm3383 (5.4× in BNAP/GRP1, 5.9× in GRP2/MP3, 5.7× in GRP3, and 6.2× in PFMP).

Greater overlap was seen with downregulated genes, as 811 genes were downregulated in all four populations compared to NSCs (Fig. 2B). Of these, the highest magnitude were 1700012B09Rik (652.4x lower in BNAP/GRP1 than NSCs, 567.3x in GRP2/MP3, 894.3x in GRP3, and 983.3x in PFMP); Ccdc113 (829.1x lower in BNAP/GRP1 than NSCs, 372.9x in GRP2/MP3, 1041.8x in GRP3, and 661.1x in PFMP); and Fam183b (1058.7.1x lower in BNAP/GRP1 than NSCs, 366.5x in GRP2/MP3, 405.3x in GRP3, and 434.7x in PFMP). As with upregulated genes, the highest overlap between populations was seen between BNAP/GRP1 and GRP2/MP3 and between GRP3 and PFMP.

### 3.6. Each neural progenitor population expresses unique upregulated and downregulated genes

Each NP population expressed unique upregulated genes compared to neural stem cells. The greatest number was seen in BNAP/GRP1 with 366 uniquely upregulated genes while GRP2/MP3, GRP3, and PFMP had 270, 100, and 206 respectively (Figure 2A). The greatest number of uniquely downregulated genes was seen in PFMPs, with 697 genes uniquely downregulated compared to NSCs. BNAP/GRP1, GRP2/MP3, and GRP3 had 160, 300, and 153 unique downregulated genes, respectively, compared to NSCs. The top ten uniquely upregulated genes by magnitude for each population are shown in Figure 2B-E (BNAP/GRP1, GRP2/MP3, GRP3, and PFMP, respectively), and the 50 highest magnitude uniquely upregulated genes were plotted in the heatmap in Figure 2F.

### 3.7. Comparison of NSC-enriched genes to available datasets

We also compared our NSC-enriched genes to other datasets from published reports on NSCs. Briefly, we selected six papers that used RNA sequencing methods to characterize adult mouse NSCs [17,20,21,29] and adolescent mouse NSCs [30,31] and that compared the NSCs to the other cells within their neurogenic niches. We compared their gene sets with the genes identified in this study as NSC-enriched (see Methods for more details). The comparative data are visualized as an Upset Plot in Figure 2F. While acknowledging that the utility of the Upset Plot is limited by the depth of sequencing and number of cells included in each dataset, it is notable that the most significant overlap between our dataset and any other dataset was for Beckervordersandforth et al., (2010), followed by Marcy et al., (2023) and then Cerebrian-Silla et al., (2021).

### 3.8. Each neural progenitor has a unique signature of up- and down-regulated transcription factors

We were interested in analyzing differences in the expression of transcription factors and identifying “hub” genes that would explain differences in gene expression between the NPs. Therefore, we compared each population’s up and downregulated genes to curated lists of mouse transcription factors (TFs) in AnimalTFDB3.0. We observed 21 downregulated transcription factors in all 4 NP populations compared to NSCs (top 10 listed in Table 1), including Foxj1, Zfp474, and Trp73. Each progenitor subtype had unique up- and downregulated TFs, listed in Tables 2 and 3. Interestingly, no transcription factor was upregulated in all 4 NPs compared to NSCs. The only gene close to reaching significance in all four progenitors was Etv1, which was significantly upregulated in GRP2/MP3, GRP3, and PFMP compared to NSCs but just failed to reach significance for BNAP/GRP1 (p adjusted = 0.056).

**Table 1.**
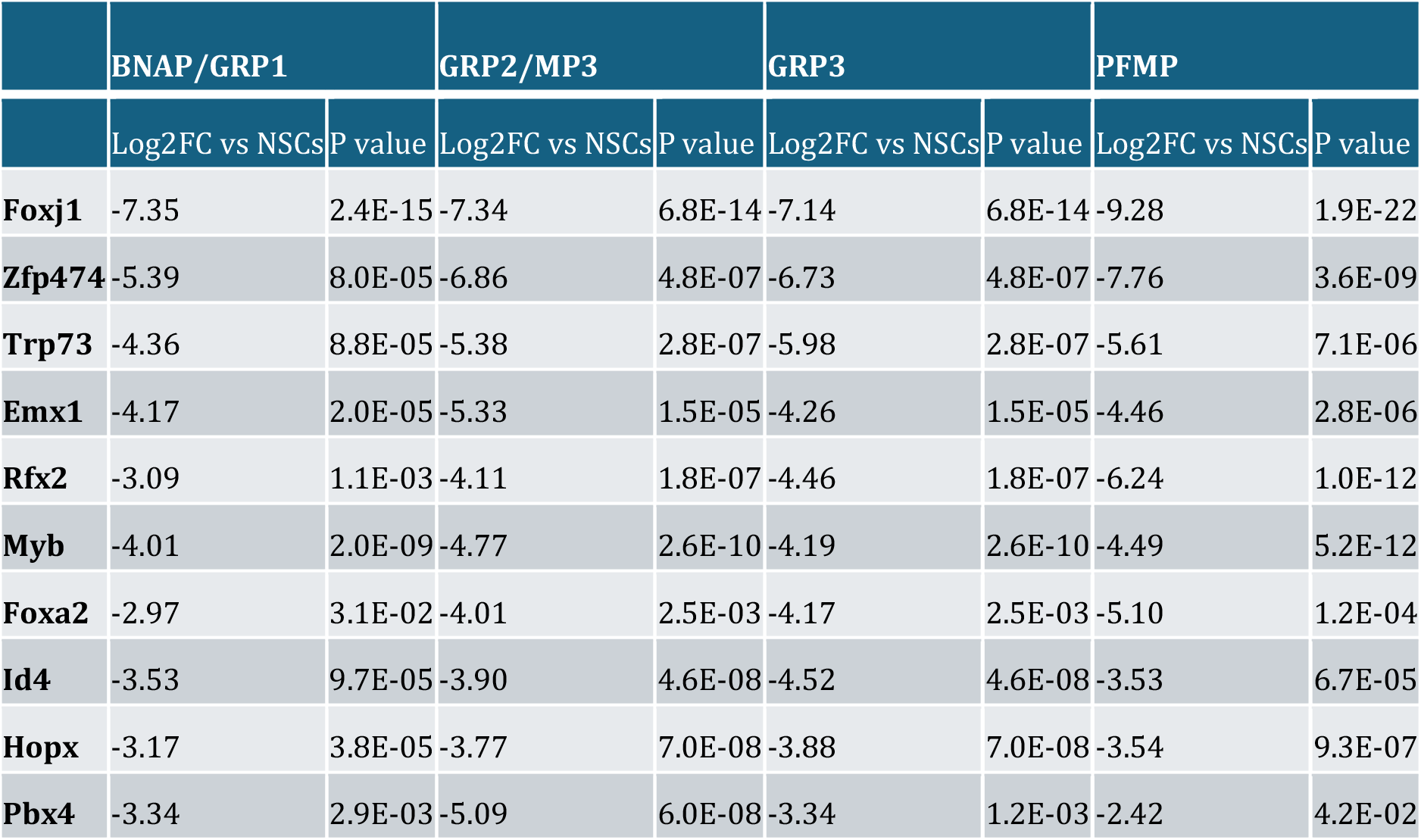
Differentially expressed genes for each subtype were compared with mouse transcription factors on AnimalTFDB3.0. Differentially expressed transcription factors were defined as transcription factors with padj < 0.05. The table lists those transcription factors that were significantly downregulated in all NP populations (BNAP/GRP1, GRP2/MP3, GRP3, PFMP) compared to neural stem cells. Included are Log2 Fold Changes vs Neural Stem cells and FDR-adjusted p-values. No genes were upregulated in all NP populations compared to NSCs.

|  | BNAP/GRP1 |  | GRP2/MP3 |  | GRP3 |  | PFMP |  |
| --- | --- | --- | --- | --- | --- | --- | --- | --- |
|  | Log2FC vs NSCs | P value | Log2FC vs NSCs | P value | Log2FC vs NSCs | P value | Log2FC vs NSCs | P value |
| **Foxj1** | -7.35 | 2.4E-15 | -7.34 | 6.8E-14 | -7.14 | 6.8E-14 | -9.28 | 1.9E-22 |
| **Zfp474** | -5.39 | 8.0E-05 | -6.86 | 4.8E-07 | -6.73 | 4.8E-07 | -7.76 | 3.6E-09 |
| **Trp73** | -4.36 | 8.8E-05 | -5.38 | 2.8E-07 | -5.98 | 2.8E-07 | -5.61 | 7.1E-06 |
| **Emx1** | -4.17 | 2.0E-05 | -5.33 | 1.5E-05 | -4.26 | 1.5E-05 | -4.46 | 2.8E-06 |
| **Rfx2** | -3.09 | 1.1E-03 | -4.11 | 1.8E-07 | -4.46 | 1.8E-07 | -6.24 | 1.0E-12 |
| **Myb** | -4.01 | 2.0E-09 | -4.77 | 2.6E-10 | -4.19 | 2.6E-10 | -4.49 | 5.2E-12 |
| **Foxa2** | -2.97 | 3.1E-02 | -4.01 | 2.5E-03 | -4.17 | 2.5E-03 | -5.10 | 1.2E-04 |
| **Id4** | -3.53 | 9.7E-05 | -3.90 | 4.6E-08 | -4.52 | 4.6E-08 | -3.53 | 6.7E-05 |
| **Hopx** | -3.17 | 3.8E-05 | -3.77 | 7.0E-08 | -3.88 | 7.0E-08 | -3.54 | 9.3E-07 |
| **Pbx4** | -3.34 | 2.9E-03 | -5.09 | 6.0E-08 | -3.34 | 1.2E-03 | -2.42 | 4.2E-02 |

**Table 2.**
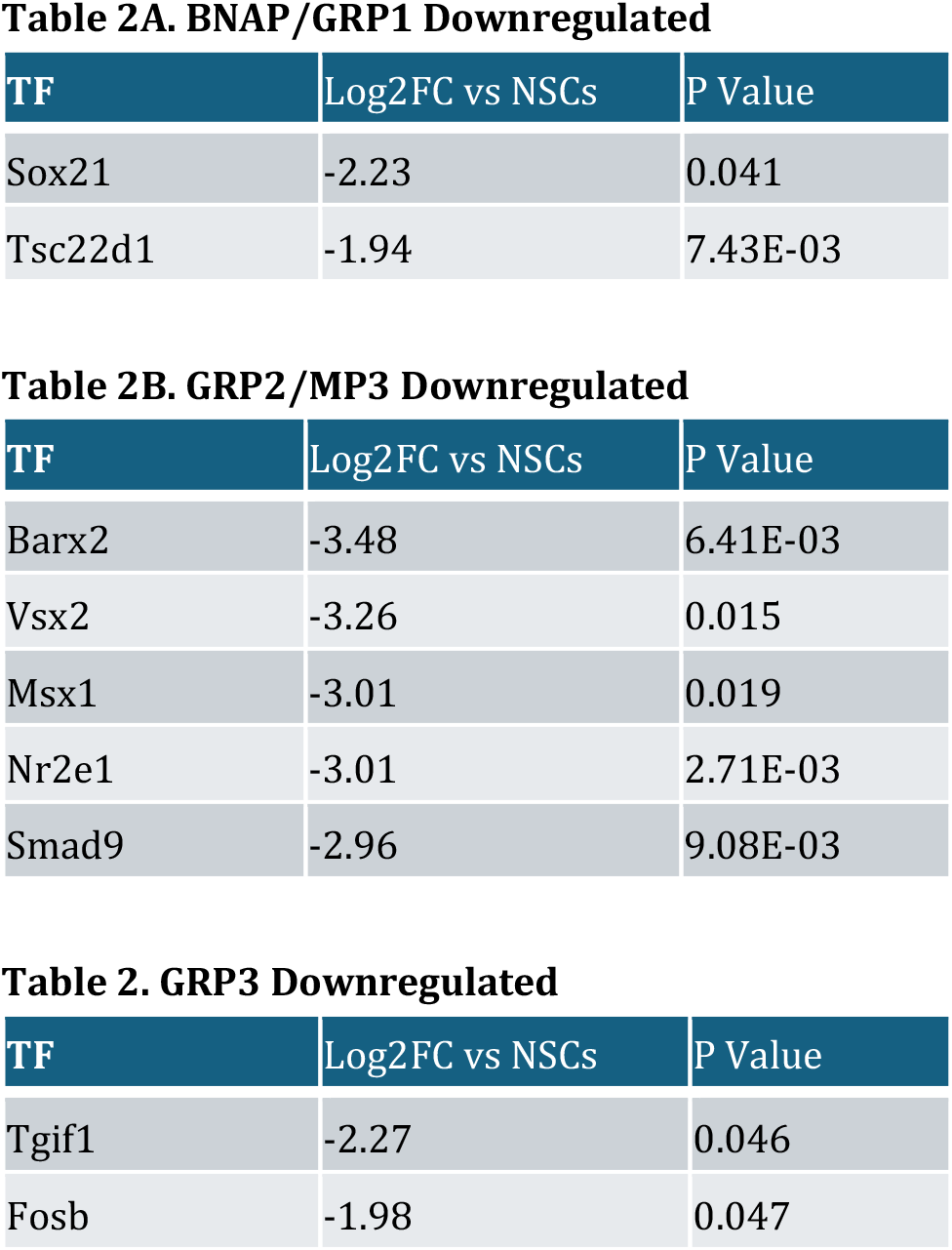
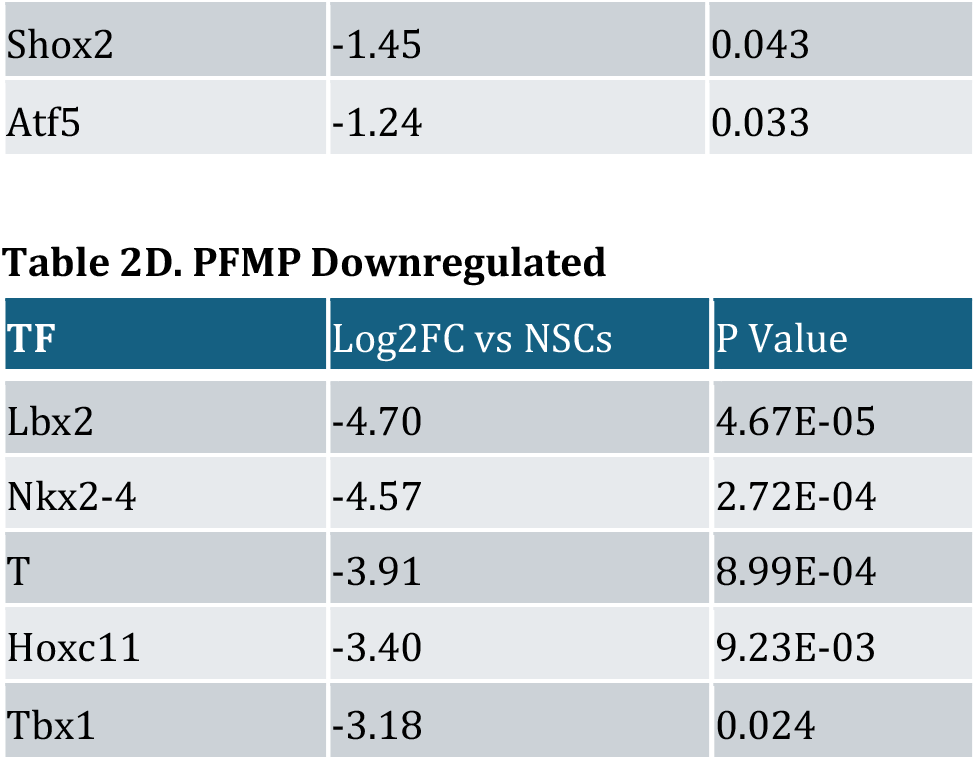
Uniquely Upregulated Genes. Differentially expressed genes for each subtype were compared with mouse transcription factors on AnimalTFDB3.0. Differentially expressed transcription factors were defined as transcription factors with padj < 0.05. The table lists those transcription factors that were uniquely upregulated in each NP population subtype: **A: BNAP/GRP1; B: GRP2/MP3; C: GRP3; and D: PFMP** compared to NSCs. Included are Log2 Fold Change vs Neural Stem cells and FDR-adjusted p-values.

**Table 3.**
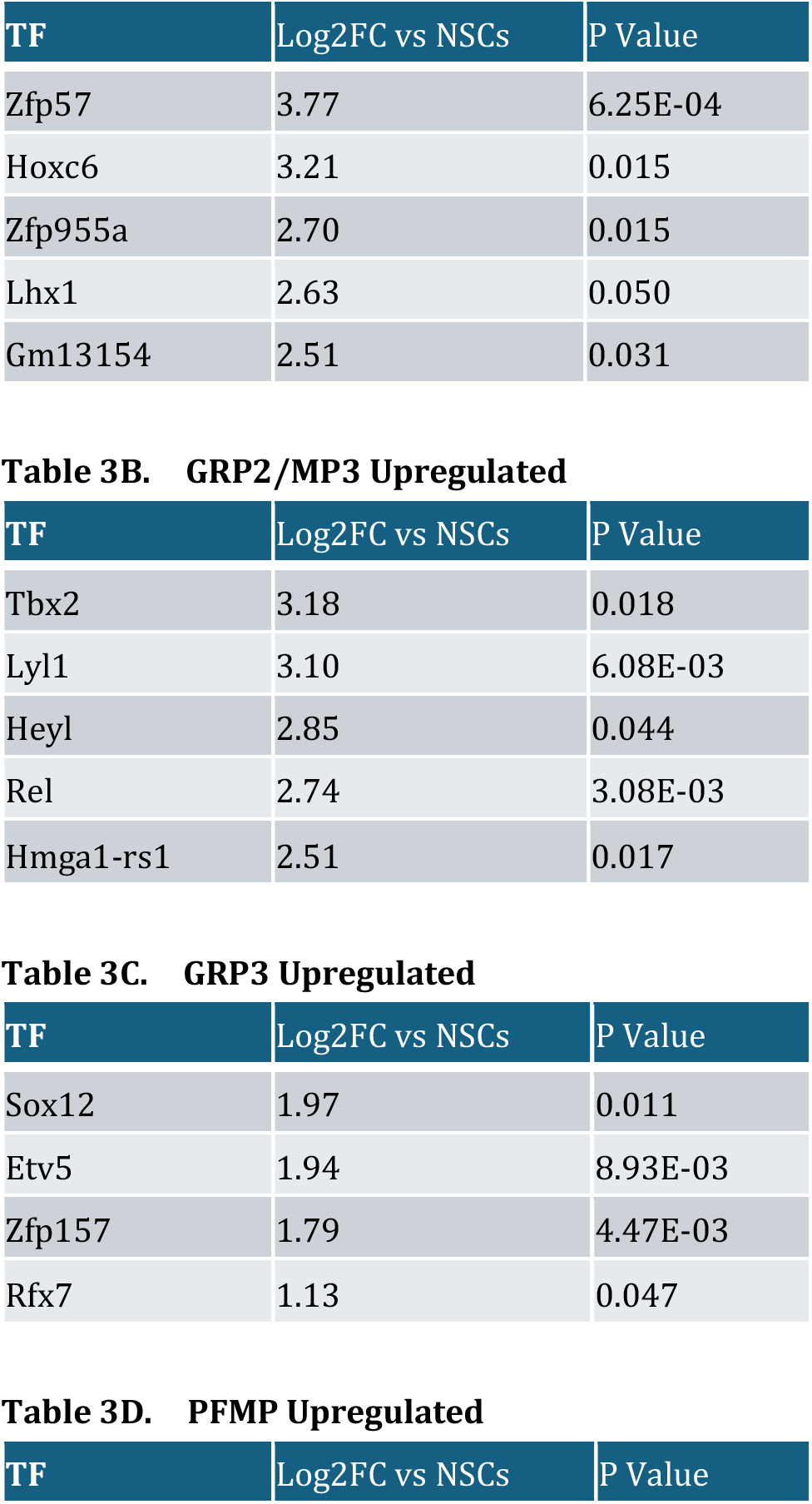
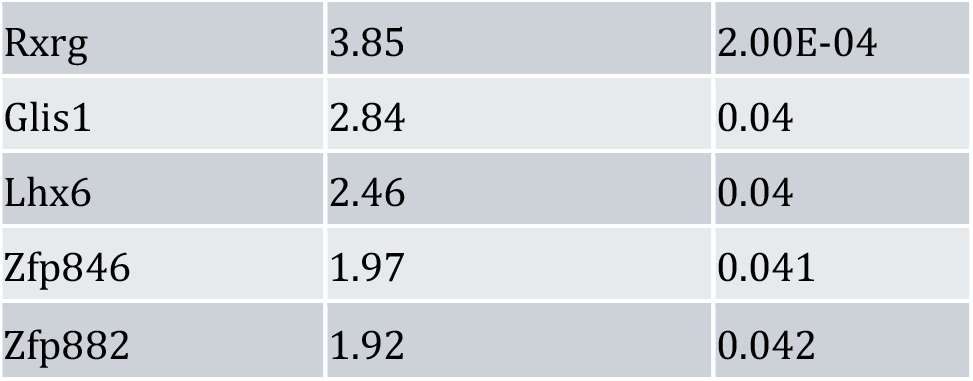
Uniquely Downregulated Genes. Differentially expressed genes for each subtype were compared with mouse transcription factors on AnimalTFDB3.0. Differentially expressed transcription factors were defined as transcription factors with padj < 0.05. The table lists those transcription factors that were uniquely downregulated in each NP population subtype: **A: BNAP/GRP1; B: GRP2/MP3; C: GRP3; and D: PFMP** compared to NSCs. Included are Log2 Fold Changes vs Neural Stem cells and FDR-adjusted p-values.

### 3.9. Neural progenitor gene expression profiles indicate unique biological functions in neurodevelopment

#### 3.9.1: Neural stem cells downregulate gene expression in cilia organization as they mature into neural progenitors

We further analyzed the expression profiles in each population using over-representation analysis, which compared our gene lists to recognized Gene Ontology gene sets (Figure S1). In each of the four NP populations, all 10 GO terms that were downregulated compared to NSCs were involved in ciliated cell function. The GO Term “Cilia Organization” (GO:0044782) was most significantly enriched with Fold Enrichment of NSCs compared to BNAP/GRP1 of 9.4-fold, 8.6 for GRP2/MP3, 9.2 for GRP3, 7.4 for PFMP) and the other 9 of the top-10 enriched GO terms had “Cilia Organization” as a parent term (Figure 3A).

**Figure 3:**
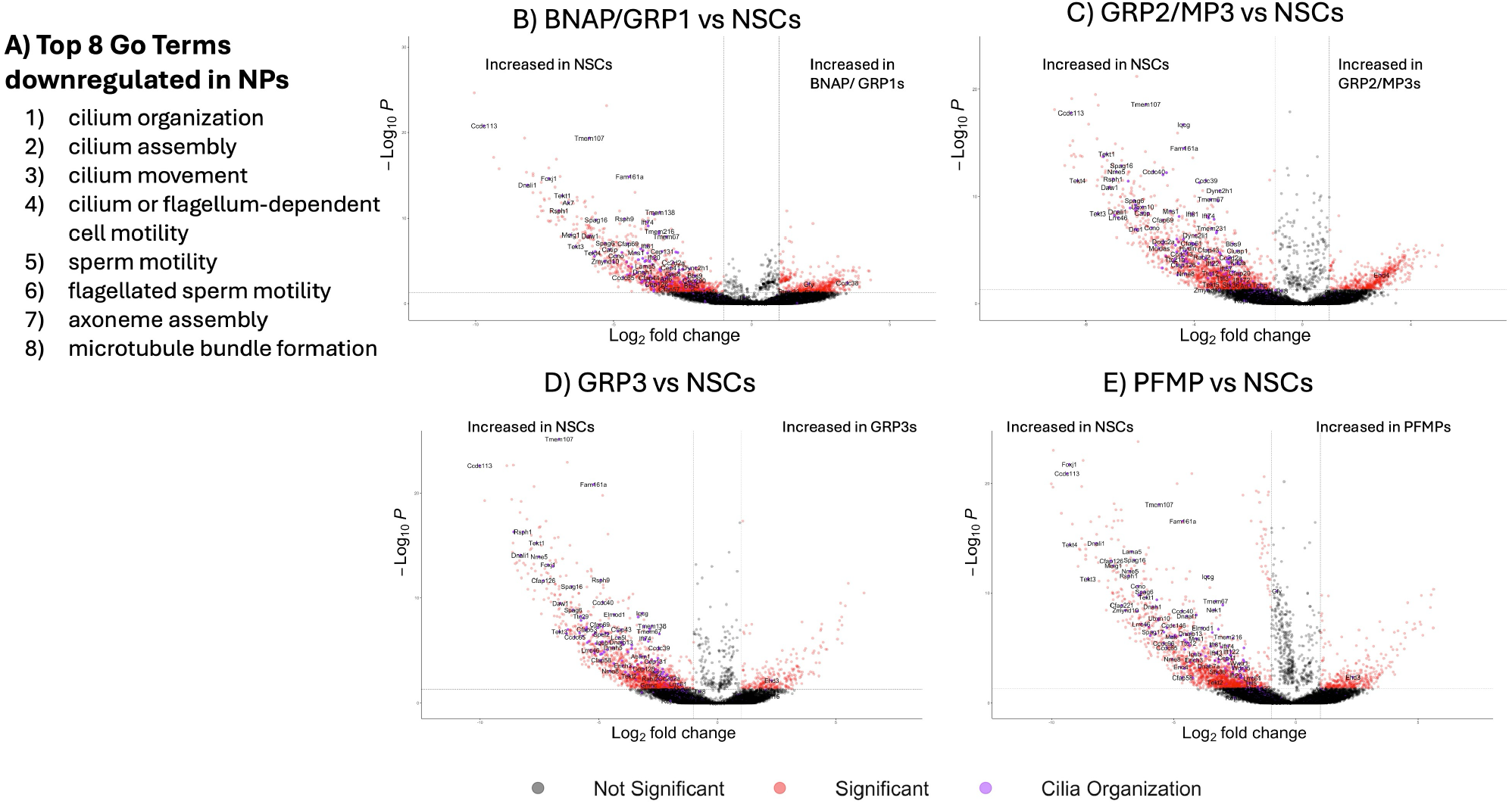
Neural Stem Cells downregulate cilia-related genes as they differentiate into NPs. **A)** Differentially expressed genes (padj < 0.05) between neural stem cells and neural progenitors were analyzed using the Gene Ontology database. Top 8 GO-terms upregulated in NSCs compared to NPs are displayed. **B-E)** Volcano plots of gene expression between each neural progenitor population and neural stem cells (**B:** BNAP/GRP1; **C:** GRP2; **D:** GRP3; **E:** PFMP). Genes in the GO Term “Cilia Organization” are colored in purple on the volcano plots and labeled.

Volcano plots of each cell type vs NSCs (Figures 3B-E for BNAP/GRP1, GRP2/MP3, GRP3, and PFMP, respectively) with cilia-related genes highlighted in purple clearly show that these genes are overrepresented in the downregulated genes in all four progenitor subtypes and make up some of the most statistically significant and high-magnitude downregulated genes. The genes with the largest change in this group were Ccdc113 (829.1-fold downregulation in BNAP/GRP1 compared to NSC, 372.9x in GRP2/MP3, 1041.7x in GRP3, and 661.1x in PFMP); Foxj1 (163.0x in BNAP/GRP1, 162.4x in GRP2/MP3, 141.4x in GRP3, and 620.5x in PFMP); and Tekt4 (54.3x in BNAP/GRP1, 315.7x in GRP2/MP3, 82.2x in GRP3, and 614.3x in PFMP).

#### 3.9.2: BNAP/GRP1 and GRP2/MP3 express many immune-related genes and pathways

The top GO terms associated with upregulated genes in BNAP/GRP1 and GRP2/MP3 emphasized immune signaling (GO:0002376, p =5.63e-4 for BNAP/GRP1 and p = 1.26e-15 for GRP2/MP3) (Figure S1, 4A,B). In addition to Gene Ontology Terms, we analyzed expression profiles using NetworkAnalyst to create upregulated protein-protein interaction networks by comparing upregulated genes in each NP population to known protein-protein interactions in the IMEx interactome. For both BNAP/GRP1 and GRP2/MP3, Sfpi1 and TNF formed central hubs of protein interactions. In GRP2/MP3, additional primary hubs included Irf8, Plcg2 and Syk (Figure 4A,B), which are uniquely upregulated in GRP2/MP3s).

**Figure 4:**
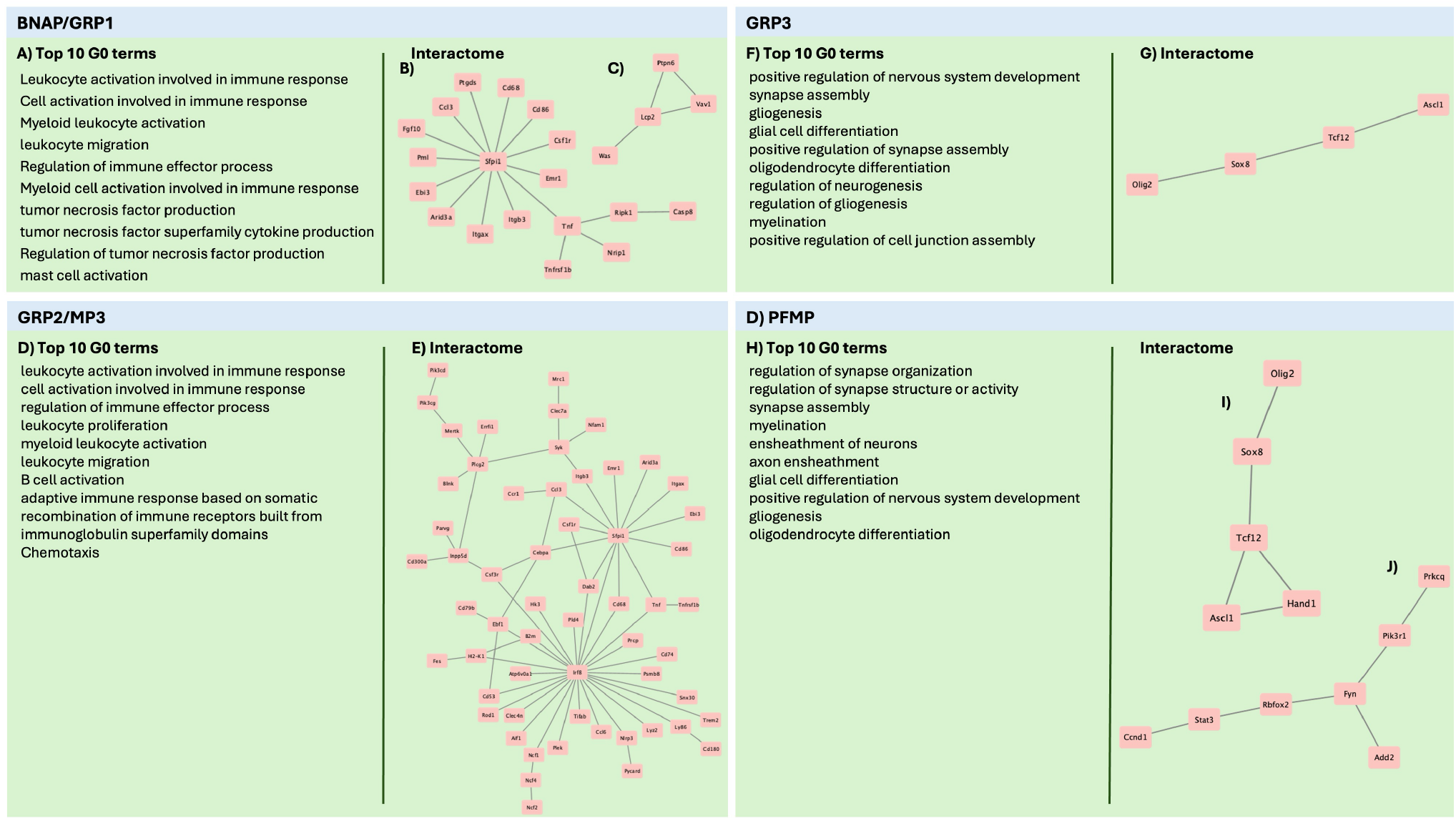
Protein-protein interactions of upregulated genes in each cell population. Genes upregulated in each population compared to neural stem cells were analyzed using Overrepresentation Analysis and compared against Gene Ontology terms. The top 10 upregulated gene ontology terms are displayed for each cell population: **A)** BNAP/GRP1; **D)** GRP2; **F)** GRP3; **H)** PFMP. Upregulated genes in each NP population compared to NSCs were analyzed using NetworkAnalyst and the IMEx Interactome. Zero-order networks with > 3 genes are displayed for each NP subtype (**B,C)** BNAP/GRP1; **E)** GRP2; **G)** GRP3; **I,J)** PFMP).

#### 3.9.3: GRP3 and PFMP upregulate genes related to neurogenesis and gliogenesis

While interactome networks were smaller in GRP3 and PMFP than in BNAP/GRP1 and GRP2/MP3, the upregulated genes and interactions highlighted are known to be involved in neurogenesis (Figure S1 and 4C,D), as reflected in the upregulated GO Terms that were related to neurogenesis and gliogenesis in both populations. The shared network (GRP3 Network and PFMP interactome Network 1; Figures 4G and 4I) include genes involved in cell fate commitment (GO:0045165, GRP3: p= 4.83e-3. PFMP: p= 5.85e-4), regulation of neurogenesis (GO:0050767, GRP3 p=2.56e-6, PFMP p=5.08e-3), and gliogenesis (GO:0042063. GRP3 p=2.81e-3, PFMP 5.21e-3). PFMPs also had an additional upregulated network (Figure 4J) involved in signaling pathways including import of proteins into the nucleus (GO:0006606, p = 0.023), cell differentiation (GO:0045595, p = 0.023), and cell proliferation (GO:0008283, p = 0.031).

### 3.10. Transcription factor analysis reveals common and unique hubs between neural progenitor subtypes

Upregulated genes were analyzed to identify transcription factors likely driving differentiation using Network Analyst software and the TRRUST database. Primary networks were pruned by including only transcription factors with Betweenness Centrality > 0.05 and results are shown in Figure 5. This analysis revealed 7 central transcription factors in BNAP/GRP1, 9 in GRP2/MP3, 6 in GRP3, and 5 in PFMP. Of these, only Sp1 was common to all populations. Twist1 was a hub transcription factor in BNAP/GRP1, GRP3 and PFMP but not in GRP2/MP3. Jun and Nfkb1 were expressed in BNAP/GRP1, GRP2/MP3, and GRP3 but not PFMP. Ets1 and Trp53 were expressed only in the BNAP/GRP1 and GRP2/MP3 networks while Ctnnb1b and Stat3 were expressed in the GRP3 and PFMP networks. E2f1 was unique to PFMPs, while GRP2/MP3s had four unique transcription factors: Gata4, Ppara, Cebpb, and Foxo1.

**Figure 5:**
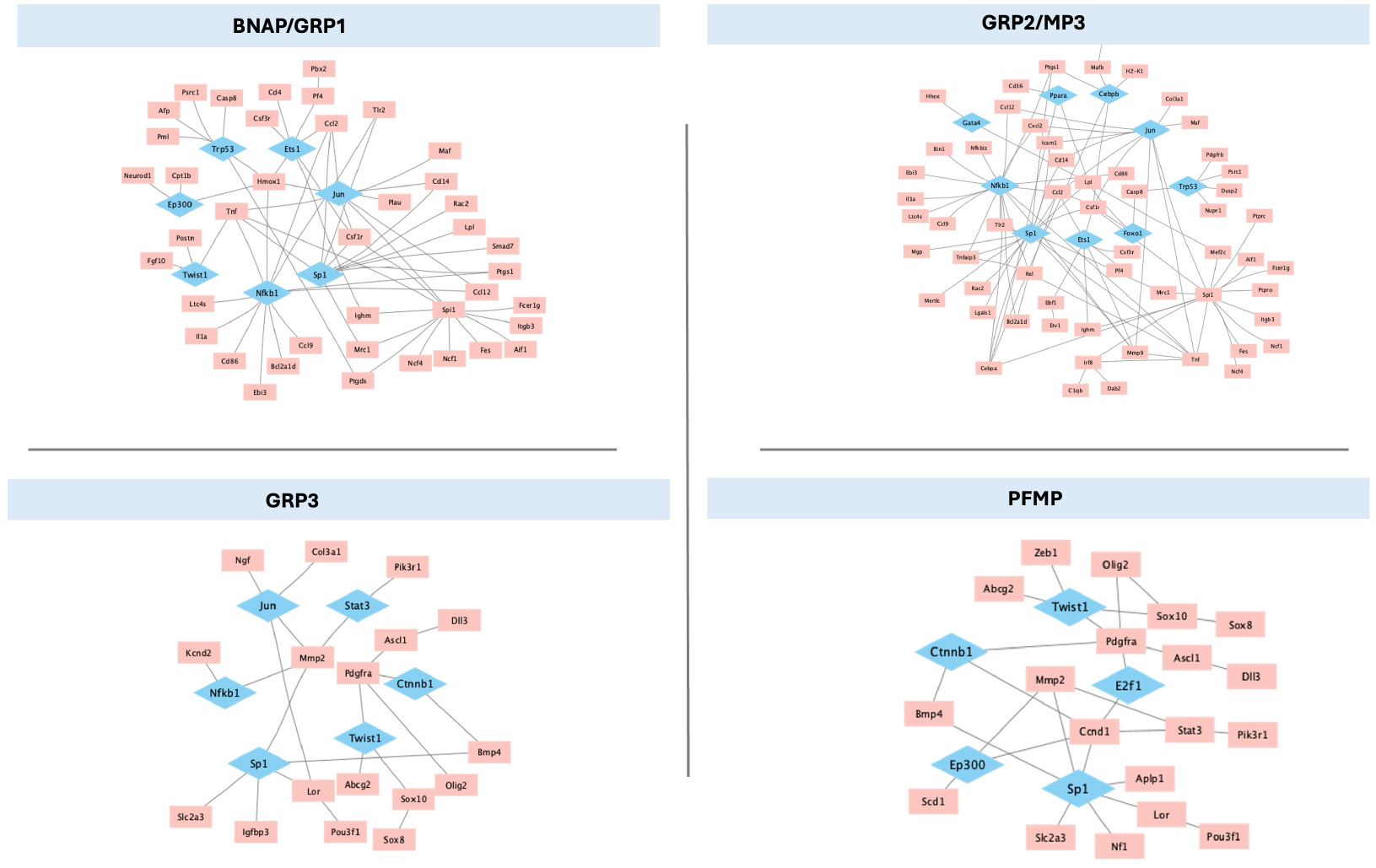
Transcription factors controlling differentiation of unique neural progenitor cell populations. Upregulated genes in each NP population compared to NSCs were analyzed using NetworkAnalyst and the TRRUST database to generate predicted gene-transcription factor interactions which were analyzed using Cytoscape. Predicted transcription factors (blue) were included if BetweenessCentrality was > 0.05. Networks are displayed for: **A)** BNAP/GRP1; **B)** GRP2; **C)** GRP3; and **D)** PFMP with upregulated genes (red) and predicted transcription factors (blue).

## 4. Discussion

In this study, we characterized four categories of SVZ resident NPs. In comparing their gene expression profiles to those of the NSCs, we have begun to discern molecular changes that occur as cells change their identity from stem cells to early progenitors. Moreover, we have identified transcription factors that are uniquely expressed by particular NPs. We have performed bioinformatic analyses to provide insights into the transcription factor interactions that are likely regulating their development, as well as the functional consequences of these differences in gene expression.

While significant progress has been made in understanding genetic signatures of NSCs, our understanding of the heterogeneity among the more abundant transit amplifying progenitors is lagging. Thus, a major gap that this study addresses is the need to provide more granular information about the transcriptional profiles of the intermediate progenitors that reside in the neonatal SVZ. Over the past decade, we have characterized seven NP subpopulations based on expression of CD133, LeX, CD140a, and NG2 [25-27]. The present study expands our understanding of the molecular differences between these different subsets of NPs by including a comprehensive transcriptomic analysis of the individual NPs. Overall, we identified 1581 genes upregulated in at least one NP compared to the NSCs (Figure 2A). Of these, 796 genes were upregulated in BNAP/GRP1 compared to NSCs; 653 in GRP2/MP3; 440 in GRP3; 527 in PFMPs (Figure 2A). We also identified 51 genes upregulated in all NP populations compared to NSCs, the most significant of which was Igf2bp3. Igf2bp3 is involved in the proliferation of NSCs and progenitors [34], which is consistent with progenitors that are rapidly dividing, in contrast with NSCs that are largely quiescent.

### 4.1. Shared TFs are expressed across NP subtypes

We specifically examined differentially expressed transcription factors (TFs) and found 21 transcription factors downregulated as NSCs become NPs. Many of these, including Foxj1 and Rfx2, function during ciliogenesis, so their loss is consistent with the loss of the NSC’s cilia during differentiation. Meanwhile, other TFs such as Emx1, Id4, and HopX are known markers of NSCs and function to maintain a quiescent state.

While no transcription factors were upregulated in all NPs, Etv1 was significant in three of the four NPs, with a very strong trend in BNAP/GRP1 (p = 0.056 adjusted). It is, therefore, possible that Etv1 activation is a hallmark of a neural stem cell’s differentiation into a neural progenitor. Etv1, also referred to as ER81, is a member of the Pea3 family of transcription factors and thus is involved in a wide variety of developmental processes [35]. It is upregulated by FGF signaling in the developing brain, as downregulation of FGF signaling significantly decreases Etv1 expression in neural development [36].

### 4.2. Neural Progenitors express unique transcription factors

We found that NP subpopulations express unique TFs in the neonatal SVZ. In BNAP/GRP1s, the top transcription factors with unique upregulation were Zfp57, Hoxac6 and Zfp955a. Zfp57 is enriched in the SVZ and rostral migratory stream of the embryo and postnatal mouse [47] and downregulated in adulthood [48]. Zfp57 is a zinc finger transcriptional repressor that maintains DNA methylation, particularly at imprinting control regions during early development; however, its function in the SVZ during neural development is not known. Hoxac6 is generally considered a marker for cervical spinal cord neurons [49], and there is no prior evidence for its expression or function in the developing brain. Hoxc6 has also been shown to bind to and regulate the expression of NCAM [50], potentially implicating it in SVZ cell migration. In gliomas, Hoxc6 is highly expressed and associated with poor prognosis. Blocking Hoxc6 in a U87 cell line caused cell cycle arrest at G0/G1 and apoptosis by increasing WIF-1, an inhibitor of Wnt activation [51]. Given the roles of Wnt in neural development [52-54], Hoxc6 may act via the Wnt pathway to promote proliferation or alter the developmental fate of BNAP/GRP1 cells. Zfp955a is increased in neurons after seizure [55], but its expression in the developing brain has not been characterized. Of particular note, Neurod1 also is uniquely upregulated in BNAP/GRP1s but unchanged in the other populations. Interestingly, in the presence of Neurod1 signaling, NG2+ cells—all of the progenitor populations in this study— are induced to form interneurons [56]. This may represent a foundational signal for neurogenesis in the BNAP group, which pushes them towards a neuronal fate.

The top transcription factors with unique upregulation in GRP2/MP3 cells were Tbx2, Lyl1, and HeyL. Tbx2 is a member of the T-box family of transcription factors, which are known to play important roles in development and in situ hybridization studies have shown that Tbx2 is expressed in the P4 SVZ [57]. While other T Box genes have been studied in the context of neural development, information on Tbx2 in this context is lacking. In the hypothalamus, Tbx2 has been shown to transcriptionally repress Shh, leading to increased differentiation and proliferation of a subset of hypothalamic NPs [58]. Tbx2 also has been implicated in various cancers as a pro-proliferative factor and may play a similar role in GRP2/MP3 [59-61]. Lyl1 is typically associated with hematopoietic stem cells and erythropoiesis [62] and also has been noted to have a role in the development of microglia [63]. While there is no known role for Lyl1 in NPs, its expression peaks in the brain at P4, the period studied in this work, and the peak of SVZ cell output [64]. HeyL is a known pro-neural transcription factor. When Jalali and colleagues grew NPs in culture, increasing Hey1 expression using a retroviral vector increased the proportion of neurons and decreased the proportion of astrocytes generated, while depleting HeyL reduced the numbers of neurons produced while promoting astrogliogenesis. Similarly, increasing Hey1 expression with a retroviral vector in E10.5 mouse embryos increased the proportions of neurons and decreased the proportion of astrocytes [65].

The top transcription factors uniquely upregulated in GRP3s are Zfp157, Sox12 and Etv5. Zfp157 is expressed in the brain, but its role in neural development has not been established [66]. Sox12 is a member of the SoxC subgroup of Sox genes along with Sox4 and Sox11. However, unlike Sox4 and Sox11, Sox12 null mice have no detectable phenotype [67]. Of note, past studies did not examine neural development or behavior in these mice, but it was noted that Sox12 null mice were grossly normal. It has been speculated that Sox12 may modulate the other members of the SoxC family, which, when induced in the neural tube of chicks, leads to expression of pro-neuronal markers [68].

Etv5 (also known as Erm) has been extensively studied for its role in neural development and has emerged as a crucial regulator of gliogenesis. In 2012, Li and colleagues showed that mice with a MEK deletion in NPs (Mek1fl/flMek2−/−NesCre) had significant impairments in gliogenesis while neurogenesis was preserved or increased [69]. The cells also showed a lack of response to CNTF (an astrocyte-stimulating signal) when differentiated in vitro. However, when they overexpressed Etv5 by electroporation into dorsal cortical radial progenitors it dramatically increased gliogenesis. Furthermore, while MEK overexpression increased gliogenesis, this effect was ameliorated by a dominant negative Etv5, indicating that MEK signaling relies on Etv5 to promote gliogenesis. Liu et al., (2019) confirmed that Etv5 knockout in NSCs in vitro decreased self-renewal and glial-associated genes while promoting neuronal genes, and that this corresponded to the production of more neurons and fewer glia [70]. They showed that this occurred due to Etv5 interacting with CoRest to bind to the promoter of Neurog2, a well-known pro-neuronal factor, and act as a repressor of Neurog2 transcription, thereby promoting a glial as opposed to a neuronal cell fate. This is particularly significant as GRP3 is the only population in this study that only produces glia, as the other two glial-restricted cell types are mixed with bipotential neuron-astrocyte or multipotential cells (BNAP/GRP1 and GRP2/MP3). Given its unique expression in GRP3s compared to all other cell types and the known correlation between ETV5 and gliogenesis, it is likely that this gene promotes the unique cell fate of the GRP3 subgroup.

The top transcription factors with unique upregulation in PFMPs are Rxrg, Glis1 and Lhx6. Retinoic acid receptors are involved in signaling throughoutneural development. RXRγ is in the RXR family of receptors, which form dimers with RARs to affect gene expression [71] and regulate various stages of neurogenesis and neural development, particularly promoting the differentiation of progenitors into neurons [72,73]. Rxrg is expressed at lower levels than Rxra and Rxrb in neurospheres [74], and, therefore, its function in development is less studied. While Rxrg mutants are viable and grossly normal, they do show impaired long-term depression in the hippocampus [75,76]. Rxrg is best studied for its role in oligodendrogliogenesis. NSCs isolated from Rxrg-null mice have impaired differentiation into oligodendrocytes and myelination. Instead of maturing, Rxrg-null cells remain in a primitive OPC-type state and they fail to exit the cell cycle [77,78].

Of the other two TFs upregulated in the PFMPs, Glis1 is a well-known transcription factor included in a panel to convert somatic cells to iPSCs [79]. It may act by stimulating proliferation genes (Mycn and Mycl) and other genes involved in development, including the Wnt and Met pathways [80]. Lhx6 is a strong predictor of interneuron fate during neural development [81,82]. In the Lhx6 null mutants, the number of interneurons (GABA+) is normal, but their distribution is altered, with cells migrating preferentially to the upper or deeper layers while absent from the middle cerebral layers. Interneuron subpopulations, including PV, SST, and CR, are also reduced, indicating a role for Lhx6 in interneuron specification [81].

### 4.3. Protein Interaction Networks and Gene Ontology

#### 4.3.1. GRP1 and GRP2s express immune signature genes

Interestingly, the over-representation analysis revealed that the GRP1 and GRP2 NPs expressed many genes for proteins involved in immune cell function. This is not surprising, since many cytokines and their receptors, complement proteins and their receptors, and MHC molecules are expressed widely throughout the brain where they regulate cell proliferation, differentiation, migration, and synaptogenesis [83-85]. Cytokines and chemokines are also well-known regulators of neurogenesis and gliogenesis [86]. The expression of receptors for immune signaling molecules likely allow the BNAP/GRP1 and GRP2/MP3 cell groups to interact with cells in their milieu, notably microglia [87]. In addition to immune pathways, GRP1 and GRP2 NPs had upregulations in pathways related to gliogenesis (p < 0.0001 for both GRP1 and GRP2) and glial migration (p < 0.0002 for both GRP1 and GRP2). However, many genes upregulated in these datasets are immune genes, including Cx3cr1, Ccl2, and Tnf.

Both GRP1 and GRP2 expressed Spfi1 (also known as Spi1) as a central hub in their protein-interaction networks, with GRP2 also having Irf8, Plcg2 and Syk as highly connected genes. Spi1, Irf8, and Syk are highly expressed in hematopoietic tissues and in microglial cells [88-90]. Spi1 also is highly expressed in the SVZ, particularly during the period studied [64]. However, data on their functions in NSCs or NPs are limited. Thus, this remains an area for further study. Plcg2, which was expressed in GRP2 cells, encodes phospholipase C gamma, a major intracellular signaling molecule. Plcg2 expression is increased by activation of the MEF2 in NSCs [91], but its role in SVZ neurogenesis or gliogenesis has not been explored. Tumor Necrosis Factor, which is highly expressed in both GRP1 and GRP2s (Figure 4), is known to be produced by NPs, where it is involved in cell survival and differentiation [92-94].

#### 4.3.2. GRP3 and PFMP express gene networks related to neurogenesis

One protein pathway that is shared between GRP3 and PFMP cell types (Figure 4G and 4I) consists of Ascl1, Olig2, Sox8 and Tcf12. All of these genes are known to be involved in neurogenesis and gliogenesis [95-97]. While Olig2 and Sox8 are typically associated with oligodendrogliogenesis [95,98], they also are involved in neurogenesis [99,100], astrogliogenesis [101-103], and the regulation of NP proliferation[104]. Interestingly, del Águila et al., (2022) [105] studied an Olig2-expressing subpopulation of cells in the SVZ in adolescent (P14) and adult mice, likely a similar population to GRP3 and PFMPs, which we have previously shown to persist in the SVZ across the lifespan [16]. Using lineage tracing in Olig2-CreER mice, del Águila and colleagues showed that Olig2+ cells generate neurons in the adult olfactory bulb and that deleting Olig2 changes the proportion of neurons generated, specifically decreasing the genesis of calretinin+ interneurons [105]. However, del Águila et al., (2022) did not examine gliogenesis, a process that continues in the SVZ throughout the lifespan, and we know from other fate mapping studies that Olig2+ cells produce glia under physiological conditions as well as some glutamatergic neurons as well [106]. Notably, the fate mapping was conducted in adult mice, so it is important to keep in mind the differences between the neonatal and adult brain, as the immature brain has much greater developmental potential.

The second network, expressed exclusively in PFMPs (Figure 4J), consists of signaling components that are less well-studied in the context of neurogenesis but likely play a role in the proliferation and migration of immature cells. For example, Ccnd1 (CyclinD1) is a known cell cycle inhibitor [107] and regulator of the balance between progenitor proliferation, migration, and neurogenesis [107,108]. Fyn regulates migration of NPs by reducing their cell-cell adhesion, potentially allowing the PFMP cells to leave the SVZ and migrate through the RMS or through the neocortex [109]. Meanwhile, the expression of Pik3r1 and Prkcq, which interact to activate PIK3CA [110], is a major growth signaling pathway throughout the body and brain. In fact, deleting PIK3CA leads to smaller brain size in vivo and fewer neurospheres in vitro [111], indicating an important role in SVZ cell division. Taken altogether, this network likely promotes the proliferation and migration of the PFMP progenitor subtype.

### 4.3. Transcription Factor Networks

Our analysis of transcription factors (Figure 5) revealed a significant overlap between the transcription factors most likely to be influencing gene expression changes in the NP populations. If this is true, it’s interesting because it indicates that the same transcription factors may have differential effects in NPs based on the cell type. The only TF present in all 4 transcription factor networks is Sp1, which is known to modulate neurogenesis by controlling both proliferation [112] and differentiation into neurons and glia [113-117]. However, the genes downstream of Sp1 differed between all four NP populations studied, with no downstream genes shared between more than two populations. As Sp1 functions largely by interacting with other transcription factors [118-120] to exert its effects, the expression and availability of interactors may drive the differential effects. Differences in downstream gene expression also may be due to differential chromatin availability for TF binding between the progenitor subtypes. This further emphasizes the point that the same stimulus, including transcription factors, can cause differential responses in each NP subtype.

### 4.4. Limitations

There are several opportunities to build upon our existing method. Specifically, this study relied on the manual dissection of the SVZ at a single time point with the sorting based on the binary expression of four antigens. This approach has its limitations. Notably, at least two mixed populations are present, with BNAP and GRP1, as well as MP3 and GRP2, sharing antigenic profiles while containing at least two subtypes of cells, according to their differentiation potentials [25]. Thus far, we have been unable to separate these subtypes using surface antigens. Furthermore, there are likely intermediate steps between the differentiation of NSCs into each NP, which we are unable to capture here, but that are surely included in the NP subtypes we did not analyze (MP1, MP2, MP4).

Another limitation of this study is that we were unable to capture the location of the NPs. Ventral/dorsal patterning is key in neural development but we were unable to discern a clear pattern of dorsal and ventral signatures [31]. Having identified genes that are uniquely expressed by each NP, our future studies should allow us to ascertain whether specific NPs are more dorsally vs. ventrally located.

There was also unavoidable contamination with cells from adjacent brain parenchyma, including pericytes, that are NG2+ [121] and likely contained within the GRP2/MP3 cell population, which can explain why we see expression of pericyte-associated genes such as Rsg5 in the GRP2/MP3 subpopulation [122].

Our information on the developmental fate of our identified progenitors is also based on in vitro analysis, and future research should aim to track subtypes of progenitors in vivo across normal development to determine their developmental fates.

## 5. Conclusions

This work details the heterogeneity of four neural progenitors that reside within the murine neonatal subventricular zone: BNAP/GRP1, GRP2/MP3, GRP3 and PFMP. We identified these cells using surface antigens and performed RNA sequencing and bioinformatic analysis to describe the differences between these progenitors and the neural stem cells. We identified differentially expressed genes and used protein-protein interaction networks and transcription factor analysis to reveal unique features of each identified cell type, which informs their biological function and that will facilitate future studies on neural development.

## Abbreviations

The following abbreviations are used in this manuscript:

NSC: Neural Stem Cell
NP: Neural Progenitor
SVZ: Subventricular Zone
FACS: Fluorescence-Activated Cell Sorting
TF: Transcription Factor
MP: Multipotential Progenitor
GRP: Glial-Restricted Progenitor
qNSC: Quiescent Neural Stem Cell
aNSC: Activated Neural Stem Cell
BNAP/GRP1: Bipotential Neuron Astrocyte Progenitor/Glial-Restricted Progenitor 1
GRP2/MP3: Glial-Restricted Progenitor 2/Multipotential Progenitor 3
GRP3: Glial Restricted Progenitor 3
PFMP PDGF-FGF: Responsive Multipotential Progenitor

## Supplementary Materials

The following supporting information can be downloaded at: https://www.mdpi.com/article/doi/s1, Figure S1: “GO enrichment across neural stem cells and progenitors.”; Table S1: Up- and down-regulated genes in each NP type compared to NSCs.

## Author Contributions

Conceptualization: SWL; funding acquisition: SWL, RZ and FJV; project supervision: SWL and FJV; experimental data acquisition: EK, RZ, FJV; Bioinformatics analyses AL and SH; writing original manuscript draft: EK, RZ and SWL. Reviewing and editing revised versions: SWL and FJV.

## Institutional Review Board Statement

All experiments were performed in accordance with Protocol #999901108 approved by the IACUC committee of Rutgers University Biomedical Health Sciences.

## Data Availability Statement

Single cell RNA-seq data will be deposited at GEO by the time of publication. All code used in this paper is available at github.com/LevisonLab/LevisonLab.

## Acknowledgments

The authors are grateful to Dr. Sukhwinder Singh and Tammy Galenkamp at the flow cytometer core facility.

