## Supplementary figures and images for "Transcriptional profiling defines unique subtypes of transit amplifying neural progenitors within the neonatal mouse subventricular zone"

### Supplemental Fig. 1

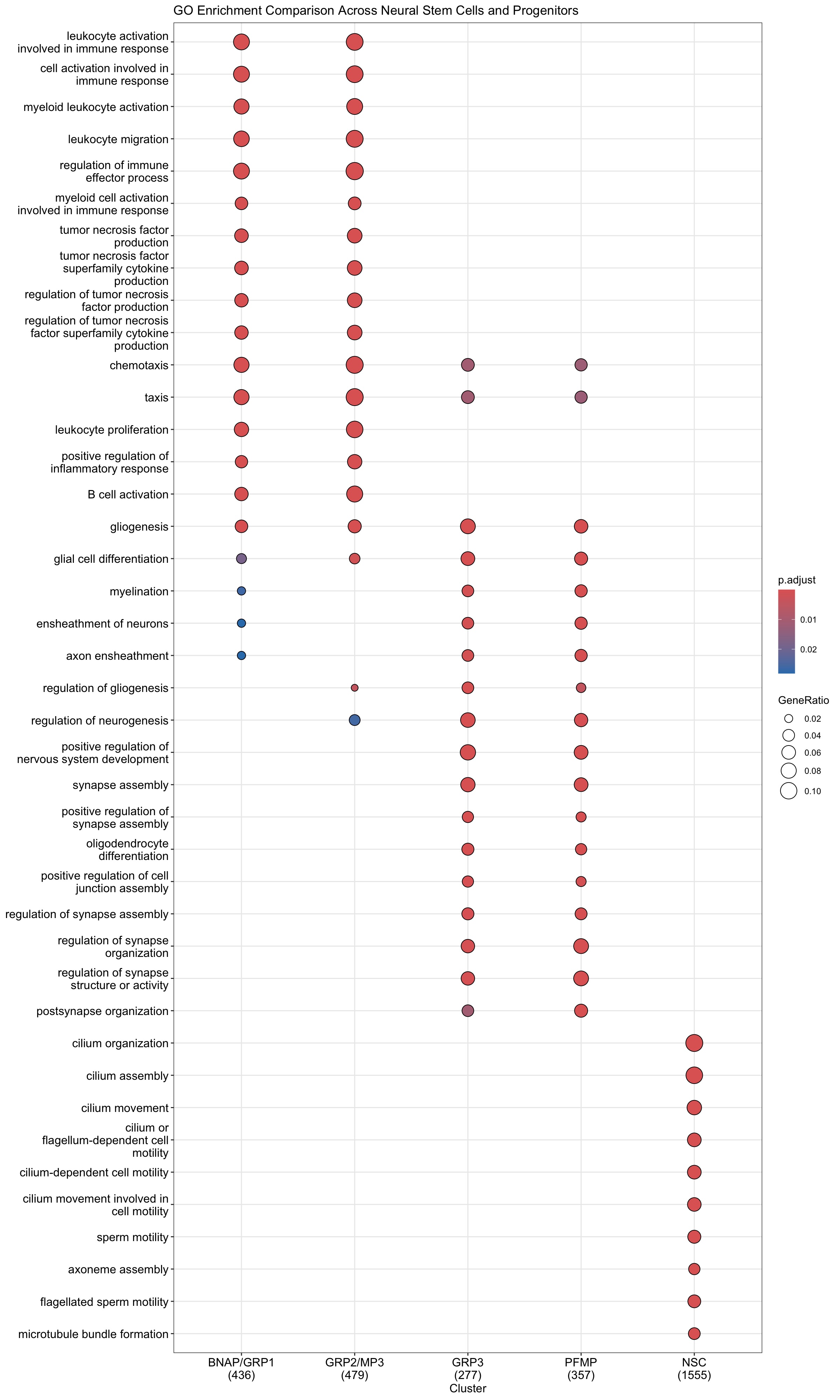
